# ROCker models for reliable detection and typing of short-read sequences carrying *mcr*, *erm*, *mph*, and *lnu* antibiotic resistance genes

**DOI:** 10.1101/2025.02.13.638121

**Authors:** Roth Conrad, Kenji Gerhardt, Konstantinos T. Konstantinidis, Amanda J. Williams-Newkirk, Andrew Huang

## Abstract

Quantitative monitoring of emerging antimicrobial resistance genes (ARGs) using short-read sequences remains challenging due to the high frequency of amino acid functional domains and motifs shared with related but functionally distinct (non-target) proteins. To facilitate ARG monitoring efforts using unassembled short-reads, we present novel ROCker models for *mcr*, *mph*, *erm*, and *lnu* ARG families as well as models for variants of special public health concern within these families including *mcr-1*, *mphA*, *ermB*, *lnuF, lnuB,* and *lnuG* genes. For this, we curated target gene sequence sets for model training and built these models using the recently updated ROCker V2 pipeline (Gerhardt et al., in review). To validate our models, we simulated reads from the whole genome of ARG-carrying isolates spanning a range of common read lengths and used them to challenge the filtering efficacy of ROCker vs. common static filtering approaches such as similarity searches using BLASTx with various e-value thresholds or hidden Markov models. ROCker models consistently showed F1 scores up to 10x higher (31% higher on average) and lower false-positive (by 30%, on average) and false-negative (by 16%, on average) rates based on 250 bp-long reads compared to alternative methods. The ROCker models and all related reference material and data are freely available through http://enve-omics.ce.gatech.edu/rocker/models, further expanding the available model collection developed previously for other genes. Their application to short-read metagenomes, metatranscriptomes, and PCR amplicon data should facilitate more accurate classification and quantification of unassembled short-read sequences for these ARG families and specific genes.

**Significance:** Antimicrobial resistance gene families encoding *erm* and *mph* genes confer resistance to the macrolide class of antimicrobials used to treat a wide range of infections. Similarly, the *mcr* gene family confers resistance to polymyxin E (colistin), a drug of last resort for many serious drug-resistant bacterial infections, and the *lnu* gene family confers resistance to lincomycin, reserved for patients allergic to penicillin or where bacteria have developed resistance to other antimicrobials. Assessing the prevalence of these genes in clinical or environmental samples and monitoring their spreading to new pathogens are thus important for quantifying the associated public health risk. However, detecting these and other resistance genes in short-read sequence data is technically challenging. Our ROCker bioinformatic pipeline achieves reliable detection and typing of broad-range target gene sequences in complex data sets, and thus contributes toward solving an important problem in ongoing surveillance efforts of antimicrobial resistance.

## Introduction

Antimicrobial resistance continues to pose a global threat to human and veterinary health care with more than 2.8 million antimicrobial-resistant infections leading to more than 35,000 deaths reported each year in the U.S. alone (1–8). Macrolides are one of the most commonly prescribed broad-spectrum antibiotics that are effective against and used to treat Gram-positive and some Gram-negative infections by *Staphylococcus*, *Streptococcus*, *Neisseria meningitis*, *Haemophilus influenzae,* in human and veterinary clinics (9). Macrolide resistance, commonly conferred by macrolide phosphotransferase (*mph*) and erythromycin resistance methylase (*erm*), can render treatment ineffective, and it can be acquired and spread through intra- and inter-specific transfer mediated via plasmids and transposons (10, 11). Another widely used antibiotic, Polymyxin E (colistin), is effective against a variety of Gram-negative bacilli and is frequently a last resort for the treatment of carbapenem-resistant *Enterobacteriaceae* infections (12). Colistin resistance is commonly conferred by the plasmid mediated spreading of mobile colistin resistance (*mcr*) genes (13). Lincosamide is another antibiotic reserved for patients allergic to penicillin or where bacteria have developed resistance to other antimicrobials, and resistance is commonly conferred by lincosamide resistance (*lnu*) genes (14, 15). As such, these resistance genes represent a major public health threat due to their ability to inactivate clinically important antimicrobials and rapidly disseminate among pathogenic microorganisms (16). Mobilization of these genes from commensal environmental organisms into pathogens may also occur since these genes are found naturally in many non-pathogenic organisms (17). Reports continue to show an increasing and rapid spread of antimicrobial resistance in natural and built environments, emphasizing the critical need for detection and monitoring tools that can assess gene functions broadly at the community level as well as more specifically at the species level.

Antimicrobial resistance gene (ARG) monitoring programs utilizing next-generation sequencing (NGS) of isolate genomes and/or metagenomes such as The National Antimicrobial Resistance Monitoring System (NARMS) and the National Action Plan for Combating Antibiotic Resistant Bacteria (CARB) could enable improved surveillance in clinical and environmental settings. However, identifying unassembled short-read sequences carrying specific gene functions remains challenging due to the unknown latent diversity of these genes in natural microbial communities as well as the frequent sharing of amino acid functional domains and motifs with related but functionally distinct (non-target) proteins (18–20). Yet, unassembled short-read sequences are the ideal candidate for ARG monitoring since they don’t require assembly, gene prediction, or annotation, which saves computational resources and avoids any errors incurred from assembly, binning, or gene prediction failure (21, 22). Additionally, unassembled short-reads can be used to robustly quantify the relative abundance of gene functions in metagenomic or metatranscriptomic type samples and thus reveal dynamics in time- or spatial-series data.

Identification of ARGs in sequence datasets typically relies on sequence similarity searches of genes predicted from assembled short-read sequences (not the short-reads themselves) against curated databases such as ResFinder (23), the Comprehensive Antibiotic Resistance Database (CARD) (24), or the Antibiotic Resistance Gene Database (ARDB) (25). Due to the continuous release of genomes and metagenomes carrying novel gene variants, these curated databases are difficult to maintain. Further, ARG sequences are frequently annotated inconsistently or erroneously in the public domains meaning that sequences remain unassigned to a family or class (26, 27). Conversely, sequences that are related but do not encode ARG functions are sometimes misannotated as ARGs. Additionally, ARG detection with assembled gene sequences does not provide a quantitative assessment of the target genes prevalence in the environment (compared to relative abundance estimates from short-read alignments with metagenomes or metatranscriptomes). For all these reasons, as well as the technical challenges mentioned above related to shared functional motifs, the tools and databases available are not intended for unassembled short-reads and suffer from unknown false positive rates (FPR) or false negative rates (FNR) when applied to short-read data. Hence, an accurate and reliable approach for detection and quantification of specific ARGs including novel and divergent variants in unassembled short-read shotgun metagenomic data sets is still needed. It is important to note that these issues are also relevant for amplicon sequencing data sets, such as those that result from PCR assays with broad-specificity primers. In particular, it remains challenging to distinguish between ARG sequences and non-ARG sequences produced in such amplicon data sets due to co-amplification of DNA from non-target genes with shared amino acid functional domains and motifs.

To address the technical limitations in detecting *mcr*, *mph*, *erm*, and *lnu* ARGs in short-read metagenomic shotgun or amplicon sequences, we built broad and specific ROCker models for each of these antimicrobial gene families (9 models in total). To evaluate model performance, we compiled testing gene sets for each model and created simulated read sets spanning a range of common read lengths (100, 150, 250, and 300 base pairs) from the total sequence of the genomes containing these genes. Instead of using fixed filtering thresholds for short-read classification across the entire length of a gene sequence—a common approach in similarity searches for identifying antimicrobial resistance gene (ARG)-carrying sequences, which may yield an unpredictable number of false positives and negatives—ROCker employs a dynamic threshold across the reference protein sequence. That is, ROCker identifies position-specific, most-discriminative bit score thresholds in sliding windows along the target protein sequences, utilizing the receiver operating characteristic (ROC) curve (28). This dynamic threshold is then called a ROCker model and can be used to efficiently and reliable filter the results of searching unknown (query) metagenomic reads against the same reference protein sequence to identity reads encoding the target protein. Hence, ROCker can better account for discriminative vs. non-discriminative domains shared by unrelated proteins, significantly reducing false positive and negative detection (28, 29). We previously reported that ROCker often shows more than a 50- fold-lower false discovery rate (FDR) than the common practice of using fixed e-values or hidden Markov model (HMM)-based searches for genes of the microbial nitrogen cycle or β- lactamase ARGs (29). Here, we report new models for ARG families not covered by our previous work and show that our new ROCker models for the *mcr*, *erm*, *mph*, and *lnu* antimicrobial gene families achieve FDR and accuracy (F1 score) comparable to our previous results.

## Results

Briefly, ROCker models are built based on a set of positive and/or negative reference sequences input as a list of UniProt IDs, which are used to retrieve the gene and corresponding genome sequences (29). Short reads are subsequently simulated from these genome sequences and mapped against the same set of positive and negative reference protein sequences. The mapping (homology search) results are then processed using the ROC curves to identify the position-specific most discriminatory thresholds and build the model. Since ROCker models are used to filter diverse metagenomic data, it is of paramount importance to train ROCker models with the most complete set of sequences available to capture the diversity of the target protein family (positive reference sequences) and close, non-target relatives (negative reference sequences), if the latter exist.

Because each of the antimicrobial resistance gene families addressed in this study consist of several subclades, we built broad ROCker models to capture the total diversity of each gene family, and then, because some specific subclades within each gene family are of high public health interest, we also built targeted models for specific subclades within each gene family. Specifically, we built and evaluated four broad models for all *mcr*, *mph*, *erm*, and *lnu* like genes, and five subclade specific models for *mcr-1*, *mphA*, *ermB*, *lnuF,* and *lnuB/G* genes.

Historically, curating the input list of positive and negative sequence targets to build and then refine a ROCker model has been a challenging and time-consuming task that could take a researcher days to weeks of effort. Recently, we developed an update to the original ROCker pipeline (ROCker V2) (Gerhardt et. al. in review) that introduces significant code and automation improvements over the original ROCker pipeline. ROCker V2 includes two modules. ROCkIn (https://github.com/rotheconrad/ROCkIn) to aid the researcher with a streamlined and partially automated process to curate the positive and/or negative target sequence sets needed to build a ROCker model, and ROCkOut (https://github.com/KGerhardt/ROCkOut) to build, refine, test and utilize the models with a more user friendly approach than the first ROCker version. We used ROCker V2 in the present study for the advantages this entails to curate and construct a total of nine training sequence sets and finished ROCker models. The models were challenged with simulated short-read datasets compiled from an additional nine complementary testing sequence sets that were used to evaluate the performance of each finished model. Specifically, for each finished model, two sets of sequences were curated. The training sequence set was used to train the finished model, and a distinct, testing sequence set was used to build the simulated short-read dataset which we utilized to test and polish the finished models. All models developed for this manuscript are freely available for use through the Environmental Microbial Genomes Laboratory website (http://enve-omics.ce.gatech.edu/rocker/models).

### Building ROCker models

To curate a high-quality input list for each ARG family or subclade mention above and build a robust corresponding ROCker model for it, we started with a set of reference seed sequences referred to as SeedSeqs. SeedSeqs may contain both positive and negative sequence targets depending on the sequence space surrounding the target gene of interest. To start, we collected the SeedSeqs for each model from NCBI’s Reference Gene Catalog, evaluating the corresponding literature and multiple sequence alignments for each gene family. Subsequently, SeedSeqs were searched against the UniProtKB database to collect the set of known gene sequences in the surrounding sequence space which returned between 5,000 −52,000 results depending on the gene family. We referred to the sequence search results as the returned sequences or RtnSeqs. Note that RtnSeqs may also contain both positive and negative (meaning non-target) sequence targets even if the SeedsSeqs contain only positive sequence targets. Next, we filtered the RtnSeqs to remove duplicate and spurious gene matches with either 100% or ≤ 50% sequence alignment and ≤ 30% sequence similarity to the SeedSeqs reducing the RtnSeqs sets to between 584 – 5,827 gene sequences depending on the gene family. We also used the UniProtKB sequence search to find a corresponding UniProt ID (100% sequence similarity) for each SeedSeq, which was retained since the required input to build ROCker models is a list of UniProt IDs. Since we have shown previously that inclusion of highly similar sequences does not improve model performance (29), and to reduce computational burden, we clustered the retained RtnSeqs at 90% amino acid sequence similarity and chose one representative for each resulting cluster to include in the training sequence set. We then selected a secondary representative from each cluster (if there was one) to include in the testing sequence set. Thus, every training gene sequence that forms a 90% amino acid sequence similarity cluster is represented in the testing sequence set by another gene of similar but not identical sequence from a different genome in the same cluster. We used the training sequence sets to build the finished ROCker models, and the testing sequence sets to simulate short-read datasets used to evaluate the finished models. For each training and testing gene set, we manually reviewed the phylogenetic clade structure, multiple sequence alignment, taxonomic classification, and gene annotation information retrieved from ROCkIn to manually classify gene sequences as either positive or negative sequence targets and input their corresponding UniProt IDs into ROCkOut. See supplemental file 1 for the full list of input sequence data and UniProt IDs for each model.

### Evaluating ROCker models

To evaluate the ROCker models, we challenged them with their corresponding simulated short-read testing dataset, and we compared the ROCker filter results to other common methods that used fixed e-value filtering approaches such as DIAMOND, BLASTx and HMMer. We also included two additional custom static filters commonly used to filter tabular BLAST results that remove alignments with ≤ 95% sequence identity or ≤ 70% sequence alignment (Custom A) and ≤ 90% sequence identity or 50% sequence alignment (Custom B). We defined sequence alignment as alignment length / query sequence length.

After alignment and filtering with the approaches outlined above, each read was identified as true positive (TP) if it passed the filter and was labeled as target, false positive (FP) if it passed the filter and was labeled as non-target, false negative (FN) if it was rejected by the filter and was labeled as target, and true negative (TN) if it was rejected by the filter and labeled as non-target. Prediction accuracy of each method was measured as the false negative rate (FNR) which identifies the failures to detect target sequences, the false-positive rate (FPR) which identifies the failures to reject non-target sequences, and the F1 score which is a summary metric that combines precision and recall. Note that the F1 score is more comprehensive but cannot distinguish different failures in the results, and therefore, we report all other metrics in addition to F1. We used the following terms and definitions to determine the quality of results: true positive (TP); false positive (FP); false negative (FN); positives (P) = TP + FP; negatives (N) = TN + FN; false negative rate (FNR) = FN/(TP + FN); false positive rate (FPR) = FP/P; sensitivity = 1 − FNR = TP/(TP + FN); specificity = 1 − FPR = TP/P; accuracy = (TP + TN)/(P + N); precision [TP/(TP + FP)], recall [TP/(TP + FN)] and F1 score [2 × precision × recall/(precision + recall)].

### Broad and specific models for *mcr* and *mcr-1* genes

We included phosphoethanolamine transferase (*eptA*) genes in our *mcr* models because plasmid-encoded (and thus horizontally transferable) *mcr* genes share their closest sequence similarity to chromosome mediated *eptA* genes. Like *mcr* genes, *eptA* can infer colistin resistance by the phosphoethanolamine transferase modification of lipid A, and it is naturally-occurring in both pathogenic and nonpathogenic organisms (30). Additionally, due to the potential of chromosomal genes to be moved to plasmids, and due to the potential rapid spreading of plasmids from nonpathogenic to pathogenic organisms, *mcr* surveillance efforts should cast a broad net to monitor for both plasmid and chromosomally encoded phosphoethanolamine transferases. Furthermore, many genomes we surveyed for this model contained both chromosomal and plasmid mediated copies of these genes and including all these copies was necessary to robust ROCker models.

We retrieved 108 *mcr* gene sequences on May 24^th^, 2023 from NCBI’s Reference Gene Catalog (https://www.ncbi.nlm.nih.gov/pathogens/refgene) identified as *mcr-1* thru *mcr-10*, and three *eptA* genes from UniProt’s Swiss-Prot database. Sequences from each labelled *mcr* allele group (1–10) formed distinct subclades. The *eptA* genes also formed two distinct subclades (Fig. S1). We selected the following 15 representatives for each subclade to use as the *mcr* reference sequence set (SeedSeqs): EPTA_ECOLI, EPTA_SALTY, EPTA_HELPY, MCR-1.1, MCR-2.1, MCR-3.1, MCR-3.8, MCR-3.17, MCR-4.1, MCR-5.1, MCR-6.1, MCR-7.1, MCR-8.1, MCR- 9.1, MCR-10.1 (Fig. S1).

The SeedSeqs were searched against the UniProtKB using the BLAST sequence search option which returned a total of 13,067 sequences (*mcr* RtnSeqs). After deduplication and filtering we retained 5,427 *mcr* RtnSeqs, and after clustering at 90% amino acid identity level we retained 1,458 representative sequences for the *mcr* training set. We then selected 473 different representative sequences for the *mcr* testing set from the same clusters (Fig. 1, Sup. File 1). There were fewer such secondary sequences than the number of clusters because many 90% amino acid identity clusters were singletons, and thus did not contain a second sequence to use. Based on our bioinformatic analysis of the sequence space during model development and refinement, we did not find the *mcr* gene function to have any verified or clear negative sequence subclades (genes with detectable sequence similarity but distinct function); that is, the closest protein family is annotated as *eptA*, which performs the same function (Figs 1, S1, S2, Sup. File 1). Therefore, we chose to include all *mcr* and *eptA* sequences from the training set as positive reference sequences to build the broad *mcr* ROCker model and all *mcr* and *eptA* sequences from the testing set as positive reference sequences to compile the simulated short-read testing dataset used to challenge the *mcr* ROCker models. That is, we did not use any negative reference sequences for the broad *mcr* model. For the *mcr-1* subclade specific model we selected only the subclade 2 sequences to be the positive reference sequences and used all the remaining sequences as negative reference sequences (Fig. 1). Subclade 2 contained sequences for *mcr-1*, *mcr-2* and *mcr-6*. Supplemental File 1 contains all UniProt IDs used, and sequence selection information for all models.

**Figure 1.**
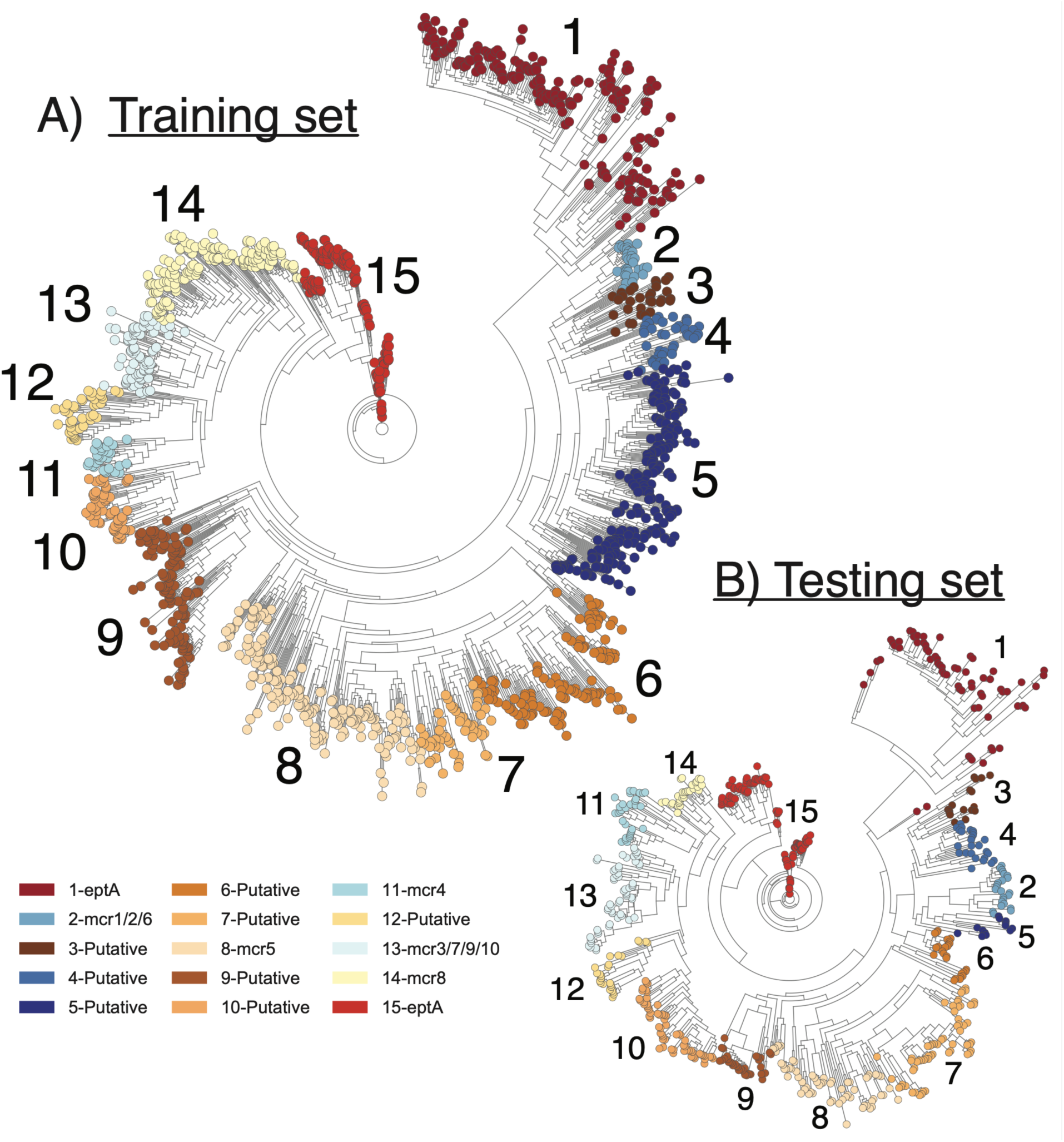
Sequence selection for *mcr* genes. Phylogenetic placement of *mcr* RtnSeqs for the *mcr* training set (A) and testing set (B). Note that there is no gene or genome overlap between the training set and testing set; different genes were selected from the 90% amino acid gene sequence clusters to match the phylogenetic diversity of the testing set with the training set to make a challenging test case for our analysis. Subclades are identified by number and corresponding color; all corresponding genes are listed with assigned subclade and annotation in Supplemental File 1. The maximum likelihood tree was constructed with IQ-Tree v2.2.2.3 using default settings. Subclades labeled as *mcr* or *eptA* contained a reference SeedSeq. Subclades labeled as putative contained only RtnSeqs from the unreviewed TrEMBL database that were annotated as *mcr*, *eptA* or phosphetholamine type functions (Sup. File 1) and showed conserved regions to our SeedSeqs in the multiple sequence alignment (Fig. S2).

The broad *mcr* ROCker model achieved clear bitscore separation between the target and non-target sequences (Fig. 2A) and an FPR of 0.053, FNR of 0.2539 and F1 score of 0.9959 (Sup. File 2) during ROCkOut’s internal model validation, indicating an excellent model fit to the training sequence set. ROCker filtering results using the simulated short-read testing dataset showed improved performance with FPR of 0.00, FNR of 0.19, and F1 of 0.90 for the 250 bp reads, especially when compared to alternative approaches (Fig. 3; Sup. File 3). More noise was observed from the shorter read lengths tested, resulting presumably from ambiguous or divergent sequence segments between the training set and testing set sequences (Fig. S3) that reduced the bitscore of these alignments (and thus, ability to discriminate between target and non-target reads) more than what was observed in the training sequence alone. Some of the HMM and static filtering methods had higher F1 scores and lower FNR than the ROCker model but at the expense of substantially larger FPR (Fig. 3).

**Figure 2.**
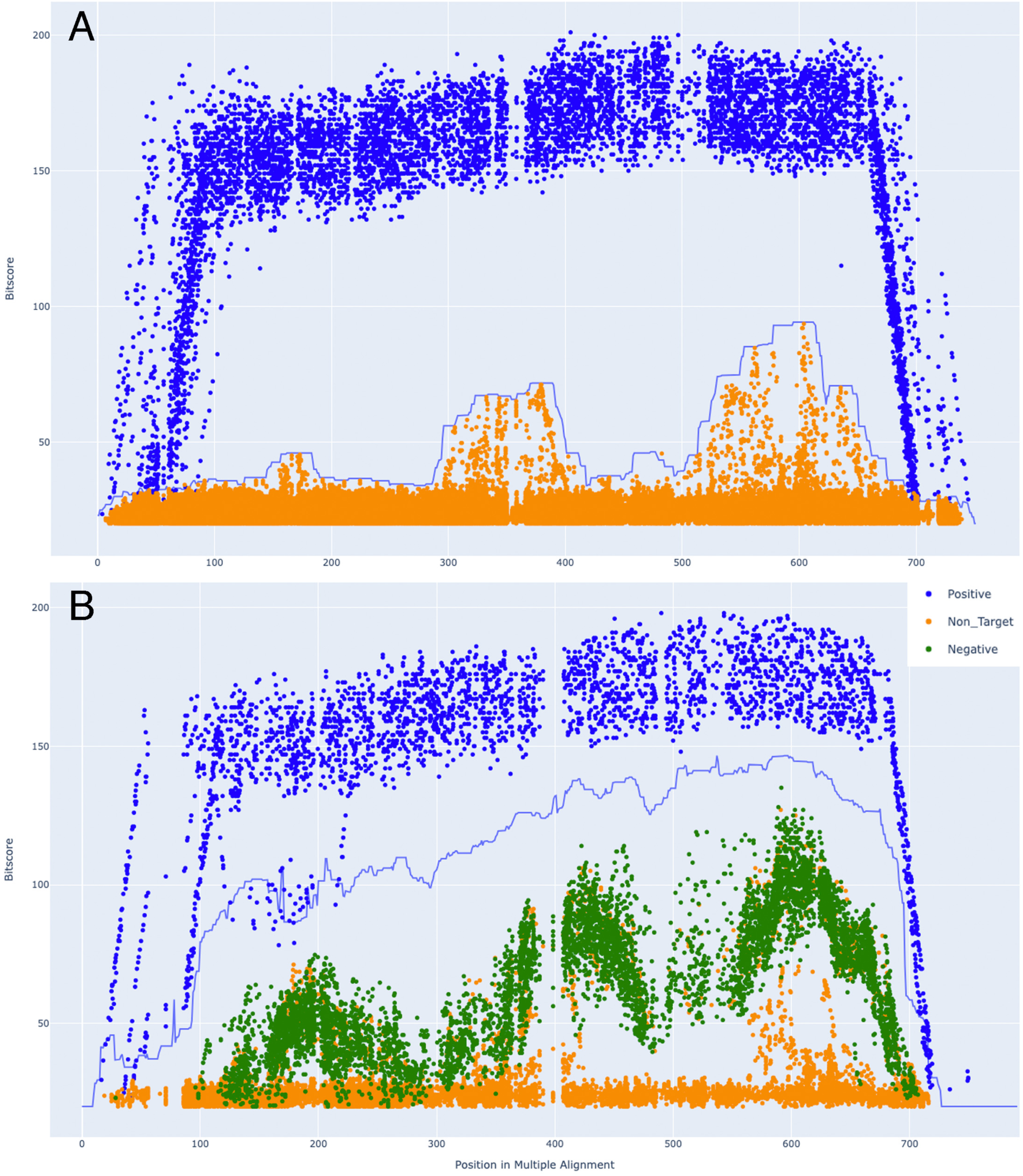
Broad and subclade specific ROCker models for *mcr* and *mcr-1*. 250-bp plots for the broad *mcr* (A) and subclade specific *mcr-1* (B) ROCker models. The graphs show the bitscore of the matching read from the Diamond BLASTx read alignments (y-axis) plotted against the amino acid position in the multiple sequence alignment (x-axis) for the corresponding model training sequence sets. The ROC curve (blue line) denotes the dynamic ROCker bitscore filtering threshold across the gene lengths between the positive reference sequence (blue) and the non-target (orange) and negative reference sequence (green) read alignments. Each dot represents the midpoint of a read alignment to the multiple sequence alignment. Note the clear separation of positive points (representing reads) above the ROC curve indicating that these models are performing well.

**Figure 3.**
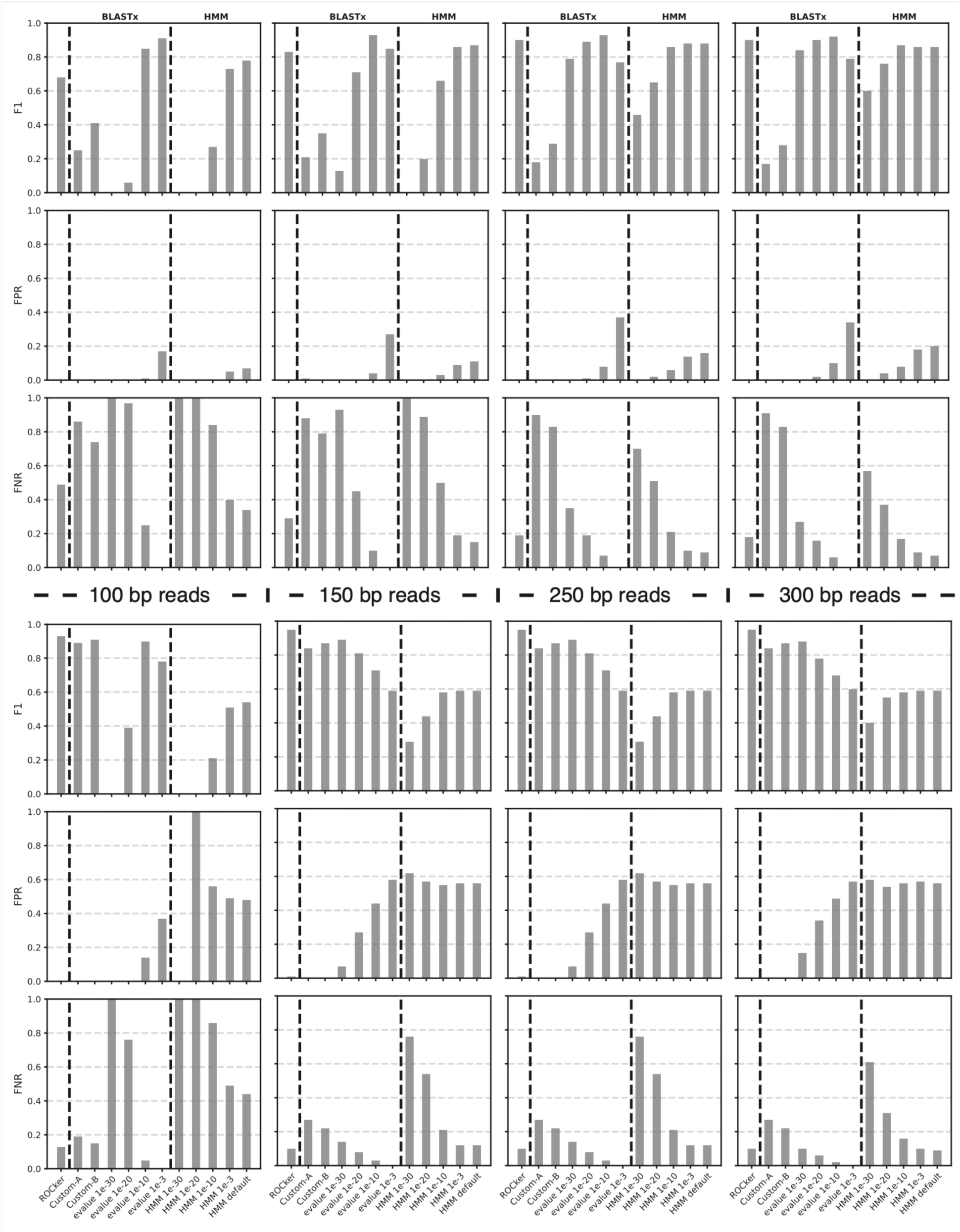
ROCker model evaluation with simulated dataset for *mcr* genes. F1, FPR, and FNR scores (y-axis) from read filtering results for the broad *mcr* (top panels) and *mcr-1* subclade specific (bottom panels) sequence sets are shown for the ROCker models and several additional commonly used static filtering thresholds for BLASTx style read mapping results or HMM search results (x-axis) using the specified e-value or a two-step filter removing matches ≤ 95% sequence identity or ≤ 70% sequence alignment (denoted as Custom A on x-axes) or ≤ 90% sequence identity or 50% sequence alignment (denoted as Custom B on x-axes). The simulated reads from the testing set were aligned to the database of training set sequences using DIAMOND BLASTx v2.0.15 with settings “--very-sensitive --unal 0” or Hmmer v3.3.2 hmmsearch program with default settings. Note the superior performance of the ROCker model (higher F1 and lower FPR and FNR) across all read lengths (by plot column) for the *mcr-1* subclade specific sequence set.

The subclade specific *mcr-1* ROCker model achieved clear bitscore separation between the target and non-target sequences as well (Fig. 2B) with an FPR of 0.0373, FNR of 0.2874 and F1 score of 0.9975 (Sup. File 2) during ROCkOut’s internal model validation, again indicating an excellent model fit to the training sequence set. In this case, ROCker filter results from the simulated short-read testing dataset had an FPR of 0.00, FNR of 0.10, and an F1 score of 0.95 for the 250 bp reads (Fig. 3; Sup. File 3). The ROCker filter results were also the most consistent across all read lengths considered.

Phylogenetic placement of the simulated short-read testing dataset reads further validated the results of ROCker and the high frequency of TP calls: target reads were mapped to the branches expected based on the reference sequence from which the read originated (Figs. 4 and S4). There were a few FP reads in the *mcr* case, and, in most of these instances, the *mcr*-carrying (TP) and nontarget-carrying reads (FP) were placed on the same branches, rendering them challenging to distinguish based on the phylogenetic approach. Since there were only a few of them, it is likely that these FP reads are a rare case from short-read simulation where a segment of a nontarget sequence became more similar to a region or functional motif in the target gene sequence. There were many more *mcr*-carrying reads that were removed by the ROCker filter (FN). While the latter reads had a similar phylogenetic distribution to TP reads, they were also similar to TN reads and their read mapping scores fell in an intermediate range (Fig. S3). These FN reads were dispersed across the full range of the *mcr* gene (Fig. S3) and were found to contain more gaps and mismatches representing the simulated reads with greatest introduced error, which presumably explains their identification as FN by ROCker.

**Figure 4.**
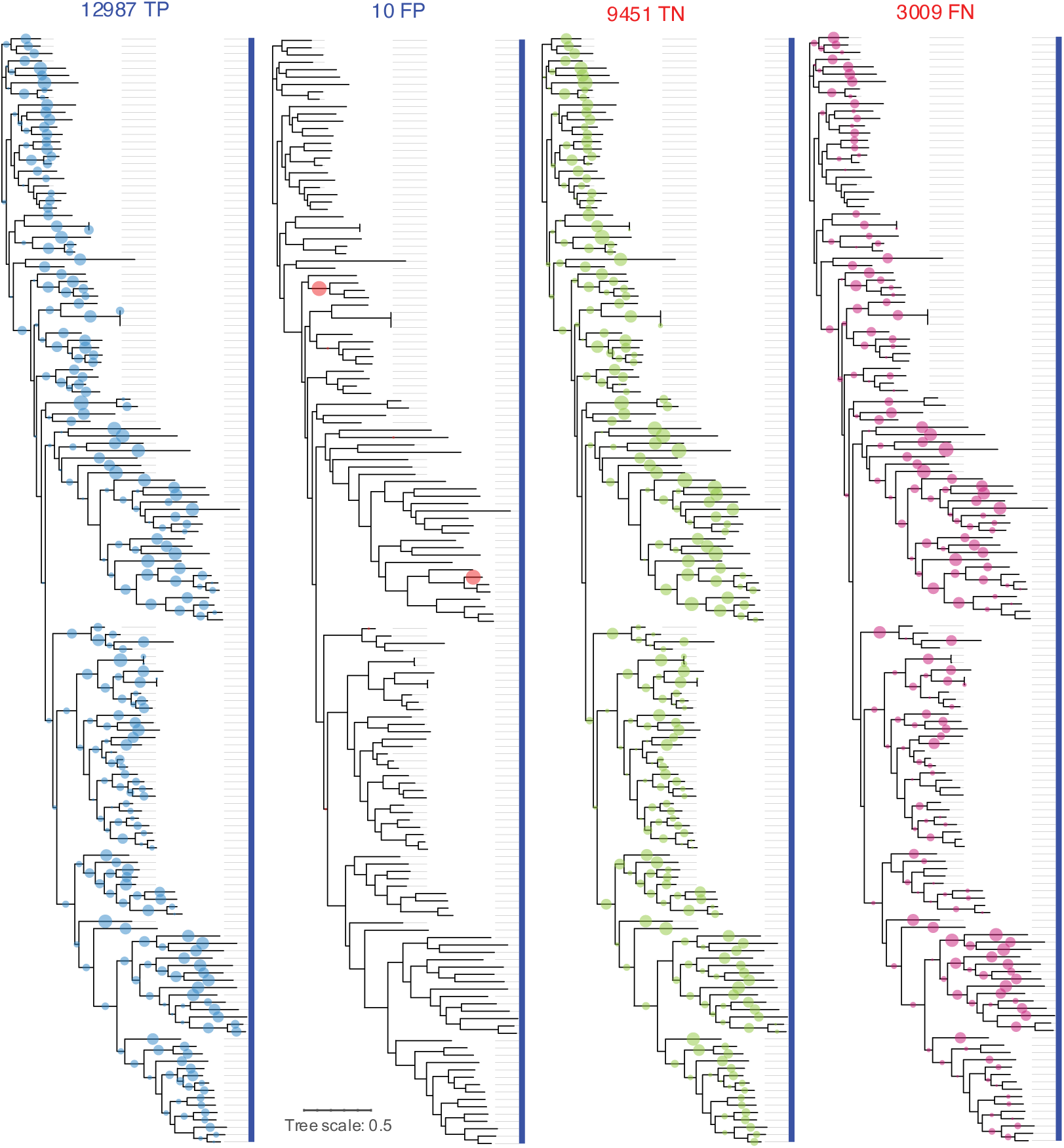
Phylogenetic read placement for the broad *mcr* model. Reads from the simulated short-read testing dataset were labeled and placed into the phylogenetic tree from the broad *mcr* ROCker model using pplacer. The size of the circle indicates number of reads assigned to each branch and the total number of reads for each category (TP, FP, TN, and FN) are listed at the top. All genes included in the broad *mcr* model were positive targets indicated by the single blue bar to the right of each tree. All four trees are the same, the short-reads mapped to the tree differ in each as indicated by the color of the circles and the label at the top of the tree.

To further evaluate the broad *mcr* model, we retrieved six random genomes, identified to contain one or more *mcr* genes, and their corresponding short read sequences from the SRA (Sup. File 4). To label the short reads from this dataset we mapped the reads to the genomes and labeled the reads that mapped to the regions of the genomes containing the *mcr* and *eptA* genes as targets. We then combined the short reads from all six genomes and ran them through the broad *mcr* ROCker model for filtering. These results from real (not simulated) short read data showed improvement compared to results from our simulated short read dataset with an FPR of 0.0027, FNR of 0.0879, and F1 of 0.95, further confirming our hypothesis that the majority of FP/FN calls by ROCker were likely due to the short read simulation inserting a greater rate of INDEL’s and SNP’s compared to real short read sequencing. Additionally, we identified that many of the FN reads from our SRA data test showed good alignments with *blastn* (nucleotide level) but poor alignments with *blastx* (amin-acid level). This is likely due to real INDEL’s or SNP’s introducing frameshifts at the amino acid alignment level leading to the FN calls. The greater INDEL and SNP rate in the simulated data would therefore lead to a higher rate of FN calls in the simulated data sets.

### Additional models for mph, erm, and lnu genes

The building and testing of ROCker models for any gene class of interest follows the same general steps and approach as described above. However, each gene class presents its own unique circumstance and results that require detailed attention. To avoid redundancy, unique aspects and results for additional models presented here are reported in the supplementary material. Overall, we observed similar performance of our ROCKer models for these other genes to those reported above for *mcr*, and thus these models are ready to be deployed for identifying reads carrying fragments of the *mph*, *erm*, and *lnu* ARGs. For instance, the average rate of FPR, FNR, and F1 for the testing datasets were 0.01, 0.15, and 0.91, respectively, for these other models (Table 2).

**Table 1:**
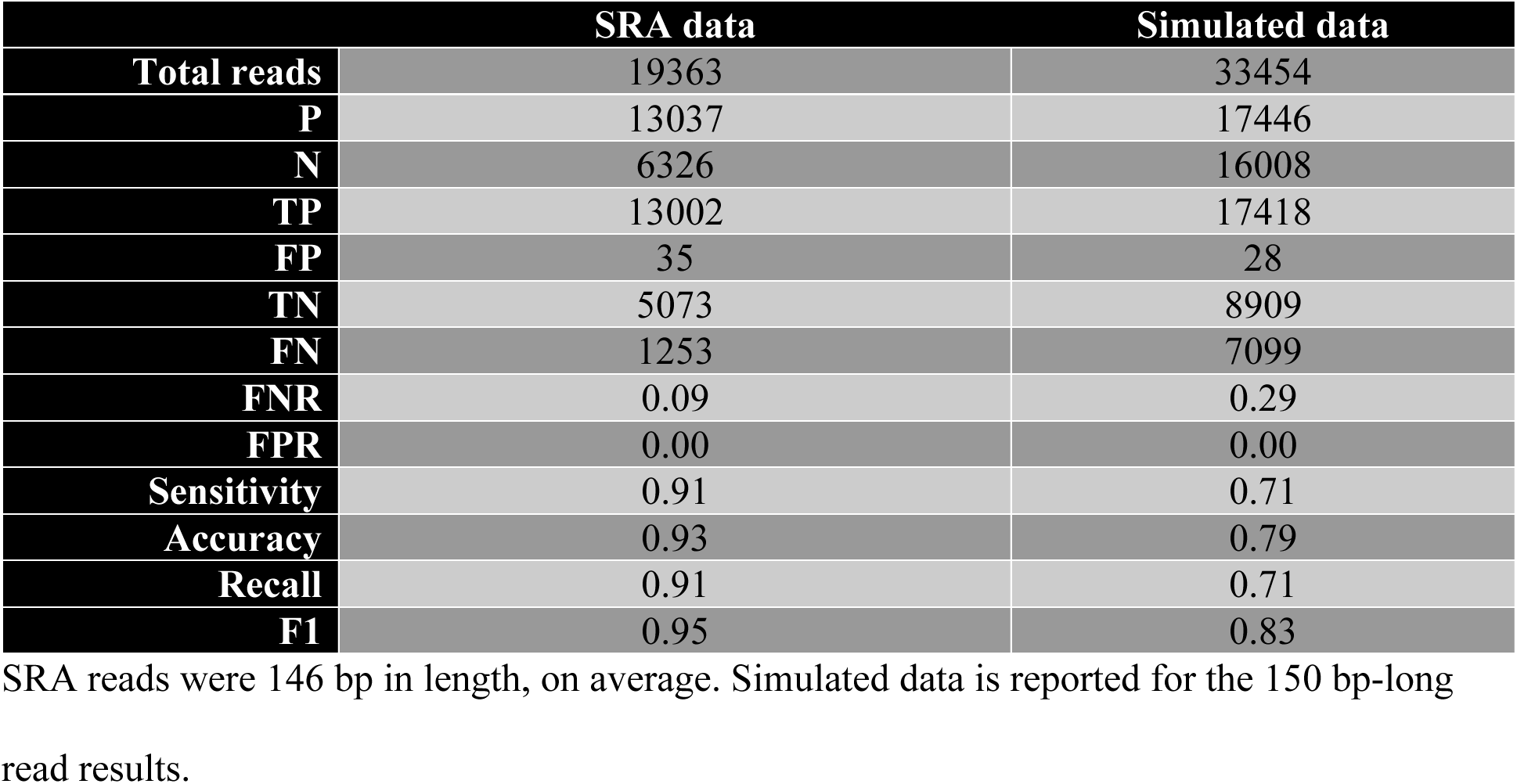
Broad mcr ROCker model filter results for SRA reads vs. simulated reads.

**Table 2:** Scoring metrics for the additional ROCker models developed here.

|  | *mph* | *mphA* | *erm* | *ermB* | *Lnu* | *lnuF* |
| --- | --- | --- | --- | --- | --- | --- |
| **FPR** | 0 | 0 | 0.04 | 0.01 | 0 | 0 |
| **FNR** | 0.1 | 0 | 0.3 | 0.33 | 0.07 | 0.1 |
| **F1** | 0.95 | 1 | 0.81 | 0.8 | 0.96 | 0.95 |

Data shown are based on the simulated 250 bp-long reads; for additional details and results for 100, 150, and 300 bp-long reads see supplemental text sections, figures S9, S16, and S23 and Sup. Files 2 and 3.

## Discussion

As macrolide resistance genes represent an increasing threat to public health, the ROCker models describe here that target some of the major genes conferring macrolide resistance determinants will facilitate the accurate quantification of these genes in high-throughput short-read datasets. Static filtering yields variable results depending on the read length and gene class considered, and it is not possible to know a priori which parameters will yield an acceptable result for a given scenario. Read filtering with ROCker models provides consistent and improved results for most cases compared to static thresholds. Furthermore, our modelling and visualization tools provide greater insight and intuition into gene class sequence diversity (e.g., which subclades to consider positive/target vs. negative sequences) as well as read mapping and filtering results.

Our analysis showed that each class of macrolide resistance gene presents its own phylogenetic and sequence classification challenges, and that there is considerable sequence diversity within each target gene class that has not been well documented yet. For example, nearly all of our identified RtnSeqs were found in the non-curated TrEMBL database (Figs. 1- S1, S5-S6, S12-S13, S19-S20; Sup. File 1). We’ve demonstrated the ability of ROCker models to perform well at broad classifications across diverse, functionally similar sequence space as well as more specific and targeted classifications distinguishing between narrower subclades within the broad class of genes. For instance, the broad *mcr* model can distinguish well across the entire sequence diversity space while the *mcr-1* subclade specific model can further differentiate these short-read sequences at finer resolution within the broad functional *mcr* gene class (i.e., identify reads representing the mcr-1 subclade to the exclusion of reads representing the other subclades), as needed for field application and/or monitoring.

While the extensive *mcr* gene diversity was more challenging for the ROCker filter to handle, e.g., relatively high frequency of FN (Figure 1), the model maintained improved results compared to alternative approaches, which improved notably with longer read length. The main source of difficulty when filtering this model was tracked to an increased number of gaps in FN read alignments comparable to gaps found in TN read alignments. It appears that a substantial fraction of these FN, if not the majority of them, is due to the read simulation step that introduced increased sequence error, leading to frameshifts errors that affected the mapping of nucleotide (reads) to amino acid (genes) sequences. Accordingly, we expect improved performance with real world metagenome datasets that are usually characterized by lower sequencing errors and artifacts due to strigent quality checking and trimming. Further, our simulated short-read testing data sets were comprised of genomes from numerous (different) species, all containing a similar functional gene. It is likely that real world datasets would contain fewer species and gene sequence diversity than our simulated datasets (i.e. lower diversity of the target sequences compared to our testing datasets); thus, increasing the likelihood of achieving improved performance. False negatives pose a more serious problem in public health contexts than false positives, where failing to identify ARGs of concern in patient samples can result in improper patient treatment and slow infection control. ROCker models performed better-in general-than alternative approaches in terms of FN (Fig. 3; Table 2), and its results could be further improved with better coverage of the sequence diversity of the target protein family. Hence, we recommend including as many reliable positive sequences as this is possible unless they are 90% or more identical to an existing positive sequence, in which case the improvement by the inclusion of such sequences is negligible -if any-.

We also observed that some gene classes, and some specific subclades within gene classes were easier to distinguish than others. For instance, the *mcr* gene class shares high sequence (and presumably evolutionary) similarity with *eptA* genes. Indeed, many genomes we evaluated contained one or more copies of both *eptA* and *mcr* genes. Likewise, the *mcr-1* subclade also contains *mcr-2* and *mcr-6,* which could not be differentiated by read mapping metrics alone. At this level, many reads may align equally well to all three *mcr* variants and more detailed analysis would be needed to locate reads mapping to specific variable regions between these genes. The ROCker models developed here allow the efficient filtering of many complete and partially matching reads to select the few reads that would be important to manually check in this case of the *mcr-1-2-6* genes.

In conclusion, our work demonstrates the ability of ROCker models to reliably detect and type short-read sequences carrying macrolide resistance genes with higher accuracy than alternative approaches using static filtering thresholds for the same purpose and type of data. Therefore, the macrolide resistance gene collection of ROCker models should facilitate their reliable detection and quantification in metagenomics, metatranscriptomics, or high-throughput PCR gene amplicon data sets from various environmental or clinical microbiomes, while avoiding false-positive calls. Furthermore, the curated ROCker models and reference *mcr* sequences available through our webserver should facilitate the development of new models for additional (phylogenetically narrow) *mcr* subclades as well as serve as a guide for the development of non-*mcr* ARGs models. Therefore, the ROCker models presented here substantially expand the toolbox for monitoring antimicrobial resistance in clinical or environmental samples and assessing public health risk.

## Methods

To develop the ROCker models presented in this manuscript, we used our new ROCkIn version 1.0 (DOI: 10.5281/zenodo.14707993) workflow with default settings (Gerhardt et al., in review) to obtain and evaluate our SeedSeqs and RtnSeqs and to assist in manual selection of the positive and/or negative sequence sets required to build ROCker models. We used ROCkOut with default settings to build, refine, and test the models (Gerhardt et al., in review). Detailed step by step notes for each model are available at the GitHub repository of the new ROCkIn/Out pipeline. (https://github.com/rotheconrad/ROCker_Macrolide_Models, DOI: 10.5281/zenodo.14708050). This repo contains everything needed to reproduce the analyses presented in this manuscript as well as all supplementary material and figures. Here we summarize the general concept of our workflow below.

### Preparations for building the ROCker Models

We used NCBI’s Pathogen Detection Reference Gene Catalog (https://www.ncbi.nlm.nih.gov/pathogens/refgene) to identify our SeedSeqs and evaluated them with the ROCkIn pipeline in order to subsequently retrieve our RtnSeqs and make our final positive and/or negative target gene lists of UniProt IDs for each model. SeedSeqs were searched against the UniProt database to retrieve RtnSeqs using EBI’s Web Services REST API Python client for NCBI BLAST+ v2.10.1 (31). The corresponding fasta sequences from the BLAST results were downloaded with EBI’s Web services Python client for WSDbfetch. The RtnSeqs were filtered, clustered, aligned, trimmed, annotated, and placed onto a template phylogenetic tree as described above and in the ROCkIn workflow. The ROCkIn workflow generates multiple sequence alignments and a data table containing functional annotation assignment and taxonomic classification that was used, in conjunction with the phylogenetic tree, to make positive and/or negative target assignments.

### Building the ROCker Models

For each model, the positive and/or negative target UniProt IDs from the training sequence sets were provided as input to ROCkOut version 1.0 modules to download, build, and refine using default settings. ROCkOut is fully automated, and these modules efficiently download all the required files, simulate the necessary data, and build the model. We followed recommended and default settings as per Gerhardt et al., in review. Essentially, model building is an iterative process that includes evaluation of the results and analysis of peculiar gene sequences to arrive at the final, well-trained model. The iterative modification details for each model are described below and in the supplemental text.

### Evaluating the ROCker Models

As part of the model building process ROCkOut simulates and labels short-read sequences for target genes and associated genomes. To compile testing datasets of known composition, we input UniProt IDs from the testing sequence sets and retrieved the labeled simulated short-reads fasta files. To evaluate the final ROCker models, we followed the same protocol used to evaluate real-world metagenomic datasets. To do this, we used ROCkOut’s align, filter, and pplacer modules with default settings providing the simulated short-read testing dataset as the input reads (-i) and the final ROCker model as the ROCker project directory (-d). The align module used DIAMOND BLASTx v2.0.15 (32) with settings “--very-sensitive --unal 0 –outfmt 6 qseqid sseqid pident length mismatch gapopen qstart qend sstart send evalue bitscore qlen slen”. The filter module used the ROCker model to filter reads and outputs files for passing and failing read alignments. The reads in simulated testing dataset reads were labeled to identify TP, FP, TN, FN. The pplacer module used PPlacer v1.1 (33) to execute phylogenetic placement of the short-reads onto the phylogenetic tree generated from the ROCker model’s target gene sequences. We used the Interactive Tree of Life (iTOL) online (34) to generate the pie chart phylogenetic tree figures.

Independently, we used Hmmer v3.3.2 (35) to build custom HMM models (hmmbuild default settings) for the same positive target gene sequences as used in the ROCker model (training sequence set) of each model. We searched our simulated short-read testing dataset against our custom HMM models (hmmsearch default settings) to generate comparative HMM alignments to the DIAMOND BLASTx alignments. We input the unfiltered DIAMOND BLASTx alignments, the ROCKer filtered DIAMOND BLASTx alignments, and the HMM alignments into a Python (36) script to read the short-read labels, apply the static e-value and custom filters, and score TP, FP, TN, FN and other associated metrics. All scripts are provided in the GitHub repository prepared for this manuscript.

## Code and data availability

All code and data details are available from:

https://github.com/rotheconrad/ROCker_Macrolide_Models

https://github.com/rotheconrad/ROCkIn

https://github.com/KGerhardt/ROCkOut

## Acknowledgments

This work was supported by funds made available from the Centers for Disease Control and Prevention Advance Molecular Detection project 223 and in part by the U.S. National Science Foundation under Award No. 2136146 and 2129823 (to K.T.K.). This project was supported in part by an appointment to the Research Participation Program at the Centers for Disease Control and Prevention administered by the Oak Ridge Institute for Science and Education through an interagency agreement between the U.S. Department of Energy and the Centers for Disease Control and Prevention.

Thanks to Jason P. Folster for assisting with manuscript review.

## Competing interests

The authors declare no competing interests.

## Author contact list

Roth E. Conrad

Kenji Gerhardt

Jason P. Folster

Amanda Williams

Andrew Huang

Konstantinos T. Konstantinidis

